## Supplementary figures and images for "Fatty acid desaturation and lipoxygenase pathways support trained immunity"

### Supplemental Figures

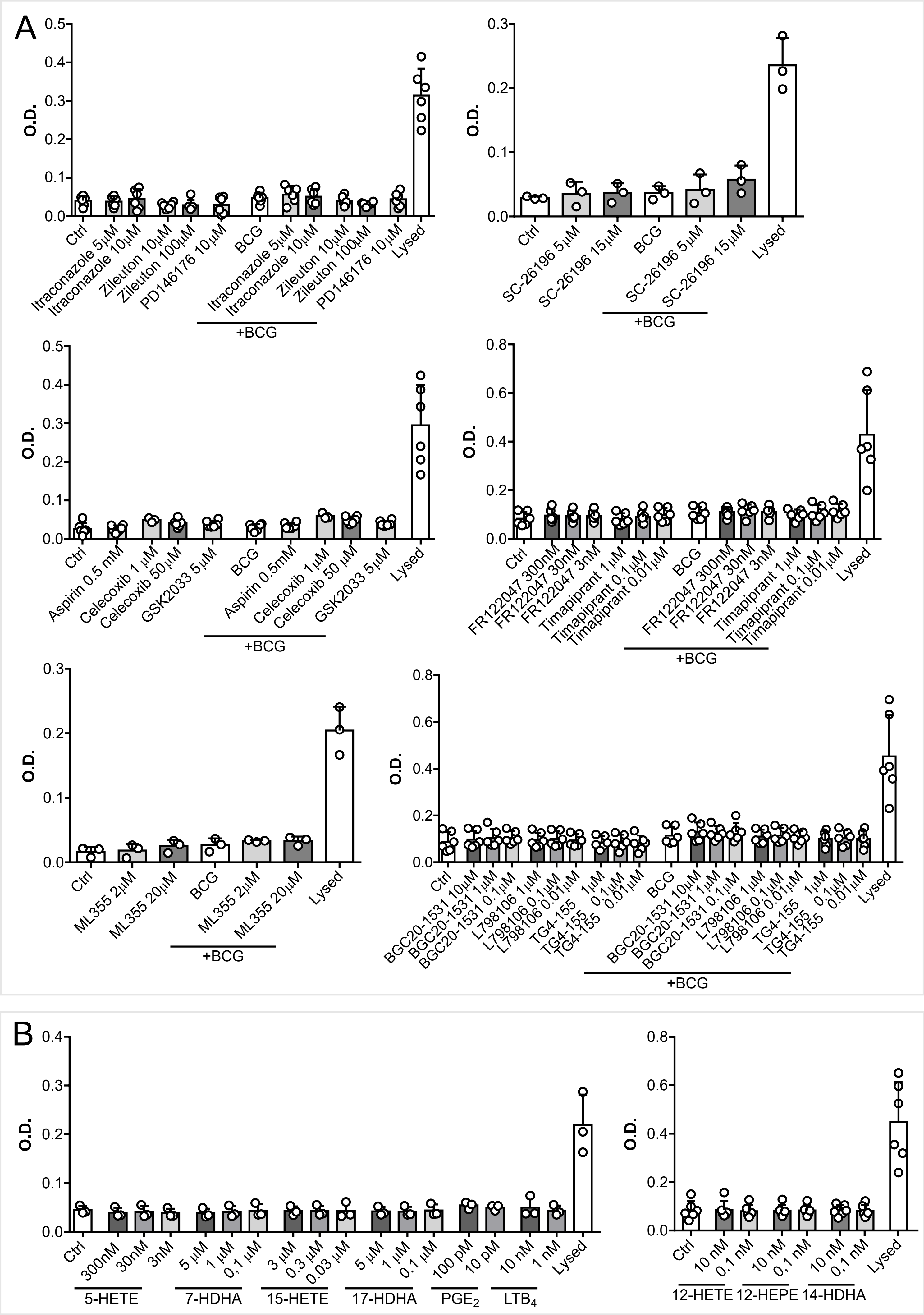

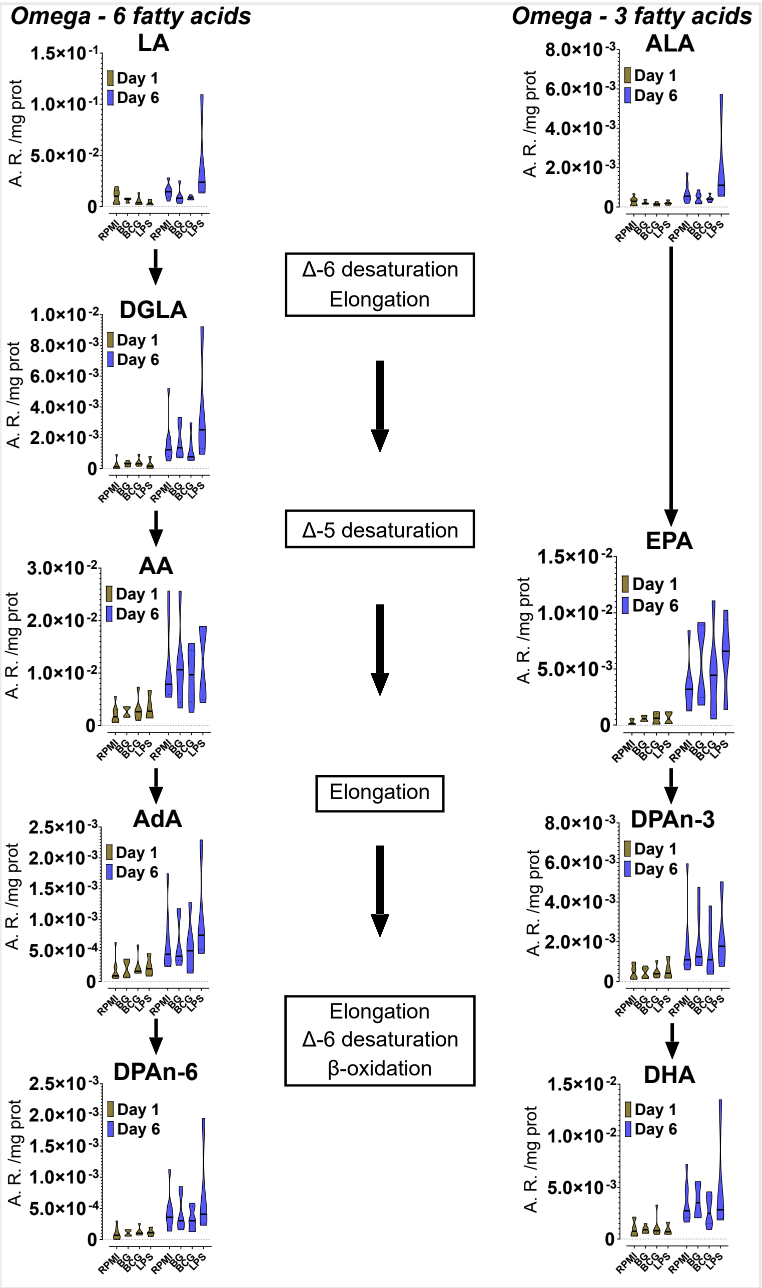

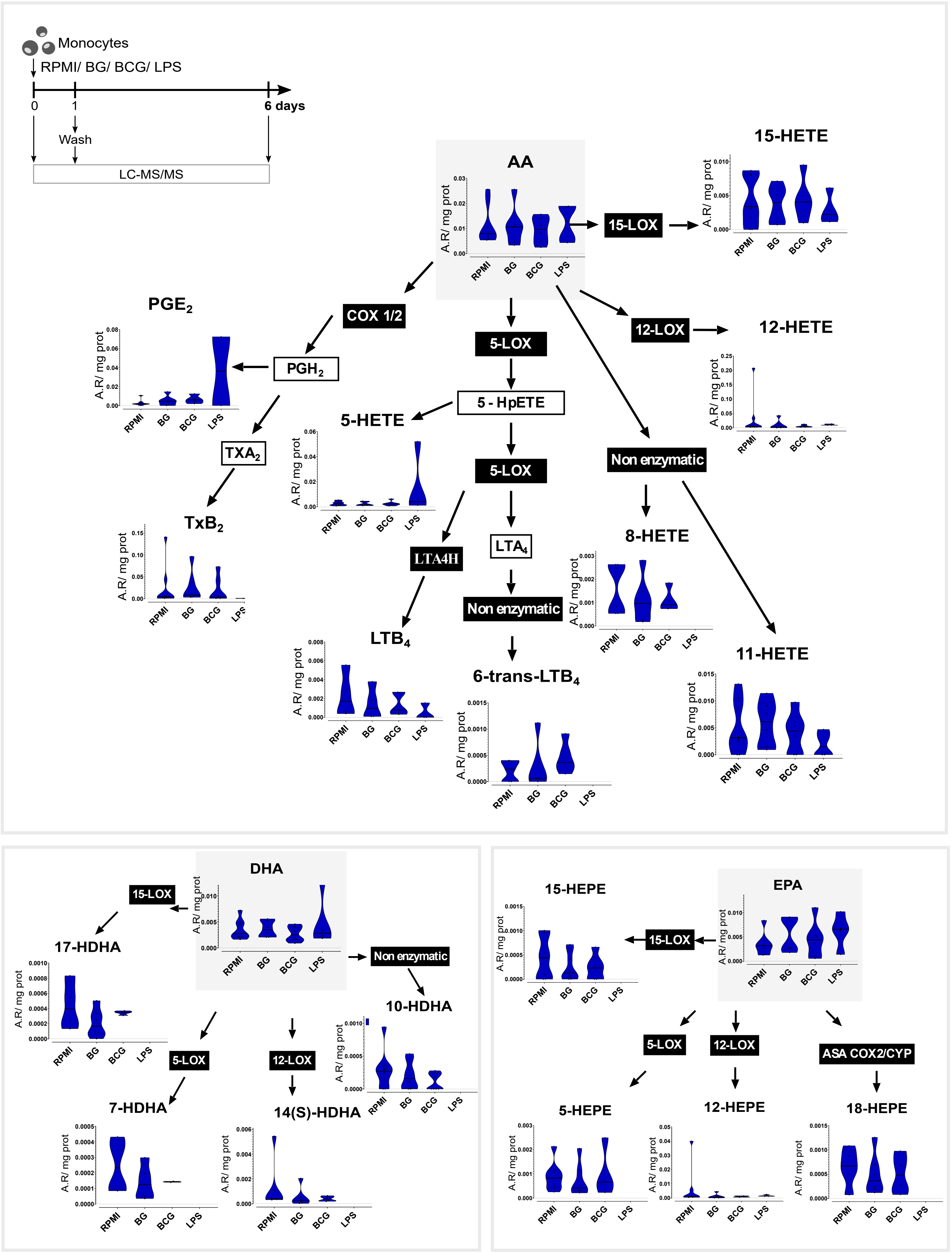

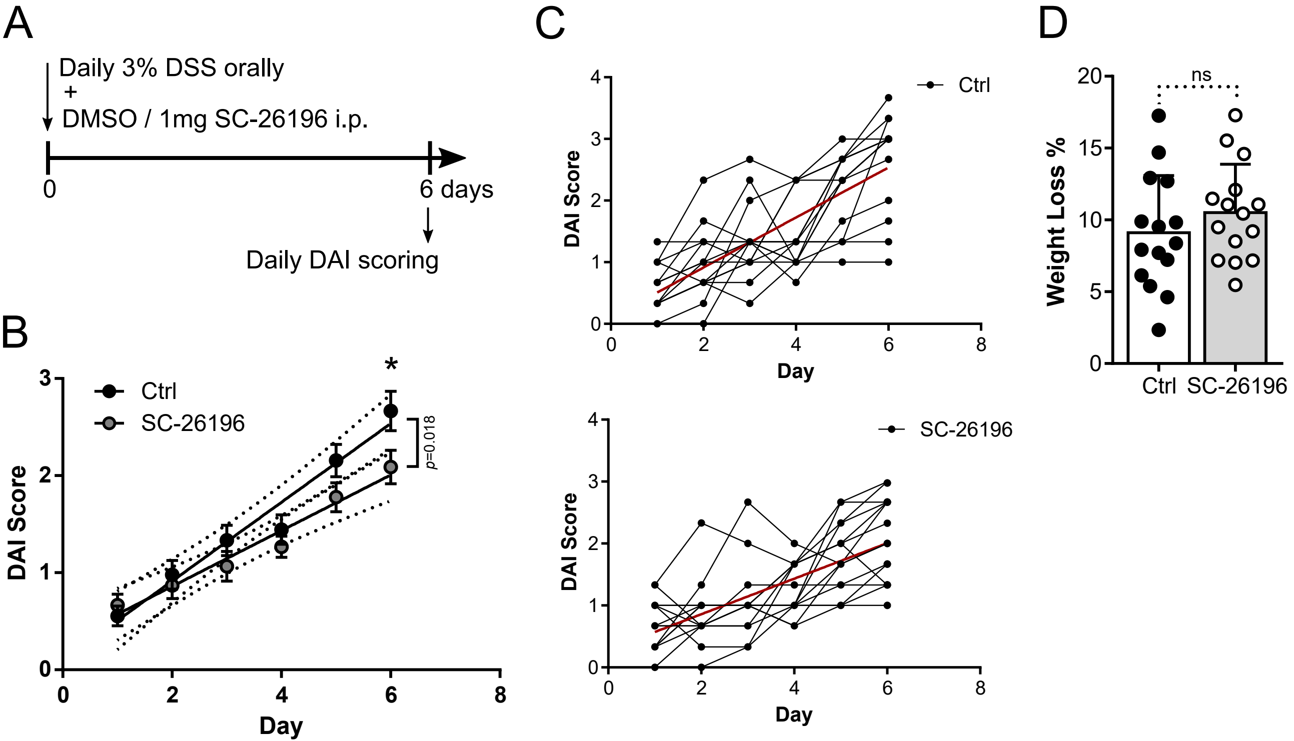

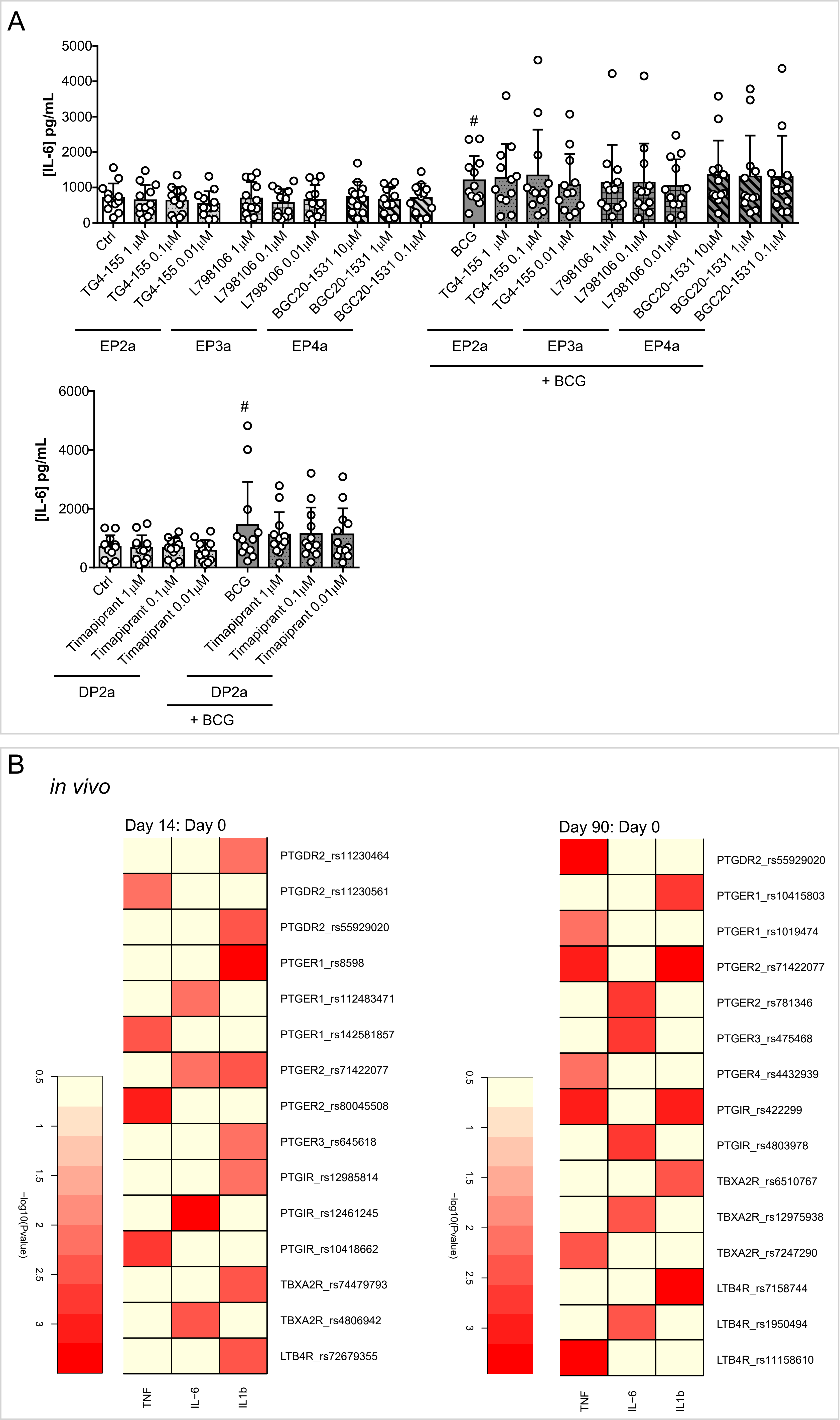

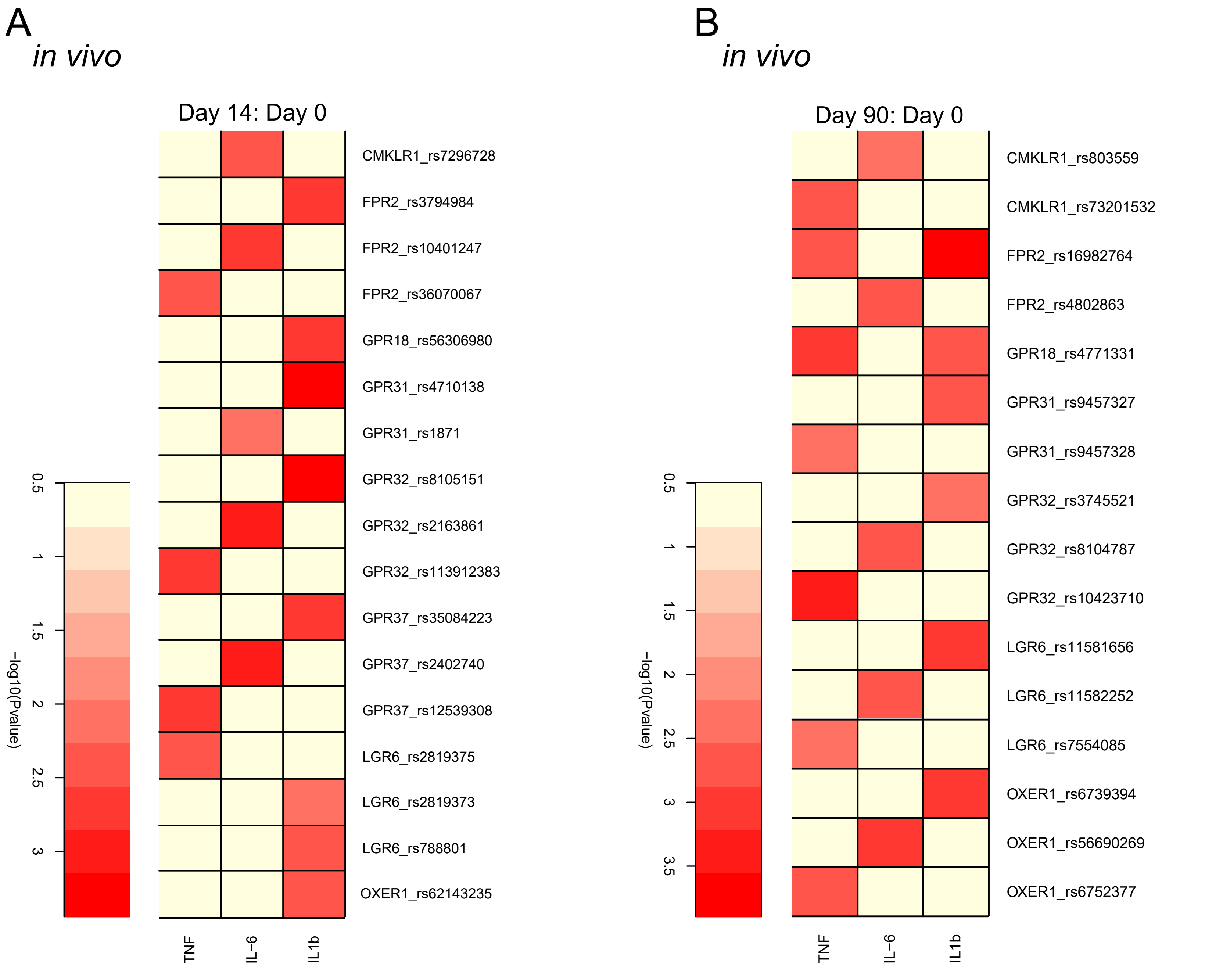

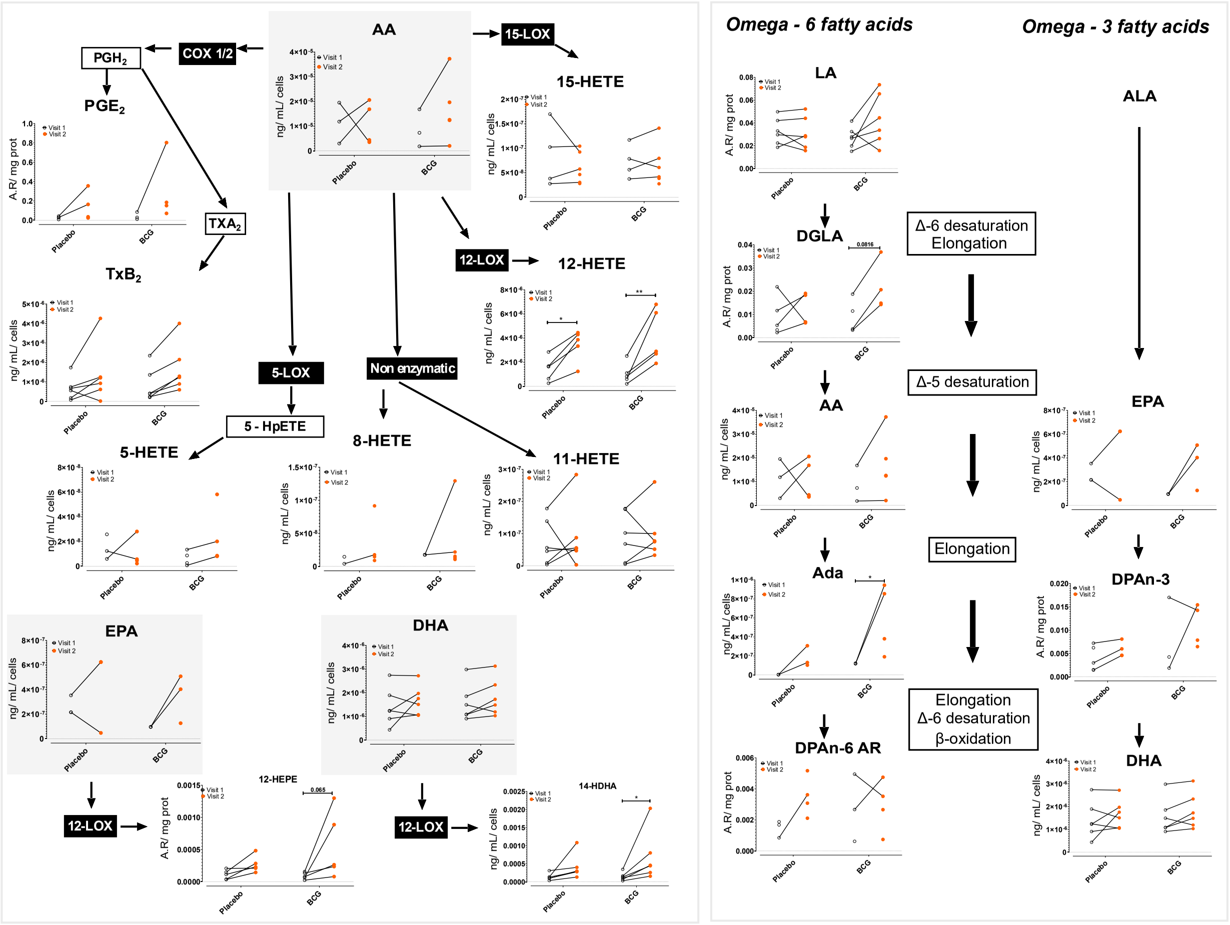
